# A Generative Neuro-Symbolic AI for Protein Sequence Design

**DOI:** 10.64898/2026.03.31.715526

**Authors:** Marianne Defresne, Delphine Dessaux, Samuel Buchet, Lucie Barthe, Liza Ammar-Khodja, Bessam Azizi, Valentin Durante, Gianluca Cioci, Simon de Givry, Alain Roussel, Luis F. Garcia-Alles, Thomas Schiex, Sophie Barbe

## Abstract

Deep learning has revolutionized computational protein design, enabling the generation of sequences that fold onto target backbones with unprecedented accuracy. However, state-of-the-art inverse folding tools largely rely on auto-regressive sampling. While powerful, this paradigm is increasingly recognized for its inability to “think ahead”, a crucial capacity to reliably create the complex, long-range inter-residue dependencies essential for most biological functions. To overcome these fundamental limitations, we introduce EffieDes, a generative neuro-symbolic AI framework that synergizes the predictive capabilities of deep learning with the logical precision of automated reasoning. EffieDes leverages deep learning to encode the target backbone’s fitness landscape into Effie— a fully decomposable probabilistic graphical model (Potts model). This landscape is then rigorously explored by an automated reasoning prover to identify sequences that simultaneously satisfy complex design constraints and optimize backbone fitness. We validated this neuro-symbolic approach through the design of orthogonal sequence pairs that adopt identical folds but exhibit selective self-assembly, as well as the design of a de novo selective nanobody with nanomolar affinity for an immune-evasive SARS-CoV-2 variant. EffieDes provides a robust architecture for precisely dissecting learned fitness landscapes, offering a new path toward proteins with highly optimized performances and sophisticated functional objectives.

## 1 Introduction

Proteins serve as the fundamental molecular machinery of life, orchestrating the vast array of biochemical processes essential for any organism’s survival. However, the repertoire of natural proteins represents only a negligible sliver of the theoretically possible sequence space. Protein design seeks to navigate this uncharted territory, employing strategies that range from the strategic modification of existing scaffolds to the de novo construction of entirely novel sequences. While rational design [1] and directed evolution [2] have historically dominated the field, they are increasingly complemented — or superseded — by computational frameworks [3–6]. By leveraging burgeoning structural and genomic databases, these digital approaches facilitate sequence-to-structure mapping while circumventing the bottlenecks of exhaustive experimental screening. Central to this evolution is the rise of deep learning (DL), which has framed protein structure prediction and design as inverse operations of a fundamental biological puzzle, ushering in a new era of predictive and generative molecular biology.

In the path from sequence to structure and eventually function, generative Protein Language Models (PLMs) [7– 9] are trained to sample sequences mimicking natural distributions. Although PLMs benefit from the vastness of known sequence space and can generate novel folds, they remain difficult to steer toward specific functional niches unexplored by nature. While PLMs have successfully produced functional enzymes [10] or GFP sequences [11], these designs typically adopt natural folds and retain significant (*>* 30%) sequence similarity to known homologs. Consequently, the design of proteins with radically new functions or architectures most often relies on a structure-based approach [3, 12]. Here, a backbone supporting the desired function is first generated [13] and subsequently “inverse-folded” to identify compatible sequences. Recently, inverse folding has been transformed by deep learning models such as GVP [14] and ProteinMPNN [15], among others [16–19]. These models leverage structural data [20] to learn potent structure-sequence mappings that frequently outperform physics-based methods, both in silico and experimentally [15].

Nevertheless, inverse folding remains a formidable computational puzzle [21]. Atomic interactions form an intricate network connecting side-chains to the backbone and to one another; a valid sequence must satisfy both local backbone compatibility and global self-consistency. To manage this complexity, most neural architectures avoid predicting all amino acids simultaneously. Even models capable of parallel prediction, like CAR-BonAra [16], eventually resort to iterative refinement. This strategy culminates in auto-regressive models such as ProteinMPNN [15], which fix amino acid identities sequentially, conditioned on previously designed positions. This approach is computationally efficient and grounded in the chain rule of probability. For a sequence of *n* amino acids *s*_1_, …, *s*_*n*_ given a backbone *B*, it writes as:

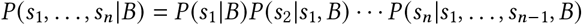

Once all conditional probabilities *P*(*s*_*i*_|*s*_*<i*_, *B*) are learned, sampling residues one-by-one from each *P* (*s*_*i*_| *s*_*<i*_, *B*)is equivalent to sampling from the joint distribution. Given that auto-regressive samplers have successfully rescued failed physics-based designs [15], they have become the standard for next-generation inverse folding tools [22]. However, auto-regressive sampling — the “Next-Token Prediction” paradigm of Large Language Models (LLMs) — suffers from a documented inability to “think ahead” [23–27]. This limitation may arise from both the learning [24] and inference processes [25]. At inference, strict adherence to the chain rule often yields average, unexceptional outputs. To access high-probability (optimized) regions, practitioners use low-temperature sampling [28] to sharpen the distribution. For ProteinMPNN, temperatures of 0.1–0.2 are routine and bring improved AlphaFold pLDDT [15, Fig. S7]. Yet, this heuristic drifts from the theoretical identity, potentially locking the generation into sub-optimal local minima. For instance, an early choice of a generic residue (e.g., alanine) might preclude the later formation of a critical, stabilizing hydrogen-bond network that a less probable initial choice (e.g., arginine) would have enabled. As residues are decoded sequentially, auto-regressive models cannot backtrack to reconsider early choices in light of emerging global constraints. In inverse folding, one can observe that very low temperature sampling eventually decreases the fraction of the highest AlphaFold pLDDT [15, Fig.S7].

This lack of look-ahead capability is well-documented in LLMs, which struggle with path-finding [25], planning [26], and reasoning tasks such as Sudoku [29]. Sudoku serves as a pertinent analogy for inverse folding: it requires selecting symbols (amino acids) subject to constraints imposed by the grid geometry (backbone) and other symbols. This analogy was first exploited by ProteinSolver [30]. Recent evidence shows that even specialized neural architectures [25, 26, 29] struggle with such problems, whereas neuro-symbolic (NeSy) approaches excel [31, 32]. NeSy models not only achieve higher accuracy but also demonstrate superior data efficiency — learning to solve Sudoku exactly from a few hundred examples, while pure deep learning models struggle even after training on millions [29]. This superiority in data-efficiency is not unexpected, as learning the full conditional probability space for auto-regressive sampling is computationally hard [24]. This efficiency implies that NeSy models may generalize better to the unseen backbones encountered in de novo protein design.

In this paper, we introduce EffieDes, a neuro-symbolic architecture explicitly designed to overcome the limitations of auto-regressive inverse folding. EffieDes separates the estimation of fitness from the sampling process. Its neural component, EffieNN, processes SE(3)-invariant backbone features [15] to predict a *joint* probability distribution represented as a pairwise Graphical Model [33], or Potts model [34]. This choice reflects the physical reality of pairwise interatomic interactions and is known to capture higher-order correlations more faithfully than variational encoders [35] or MSA-Transformers [36]. Crucially, this intermediate representation allows the use of symbolic solvers — from stochastic optimizers [37, 38] to exact automated reasoning provers [39, 40] — to solve the *Maximum A Posteriori* optimization problem. Unlike black-box neural sampling, these solvers can incorporate complex external constraints without retraining, a vital capability for functional protein design.

We evaluate EffieDes on challenging design problems where auto-regressive models should struggle. First, we target the design of a hexameric protein platform requiring orthogonal pairs of proteins that share the same fold but self-assemble *exclusively* into heteromeric pairs. This requires designing against the natural homomeric assembly favored by the backbone — a multi-state design challenge [41] where auto-regressive approaches must rely on linear combinations of logits [15], drifting further from probabilistic guarantees. Second, we apply EffieDes to the de novo design of nanobodies targeting SARS-CoV-2, requiring high-precision sequence optimization for previously unseen CDR loop structures.

## 2 Results

### 2.1 EffieNN generates deep-learned Potts models

The first core component of our neuro-symbolic design architecture, EffieNN, is a deep learning model trained on protein structure data that takes a backbone *B* as input and generates Effie^*B*^, a Potts model specific to *B* (Figure 1). Effie^*B*^ defines the score (*E s*| *B*) of any amino acid sequence *s*. This score represents the conditional probability *P* (*s* |*B*) ∝*e* ^−*E*^ (^*s* |*B*)^, where lower scores indicate sequences more suitable for the given backbone structure *B*.

**Figure 1:**
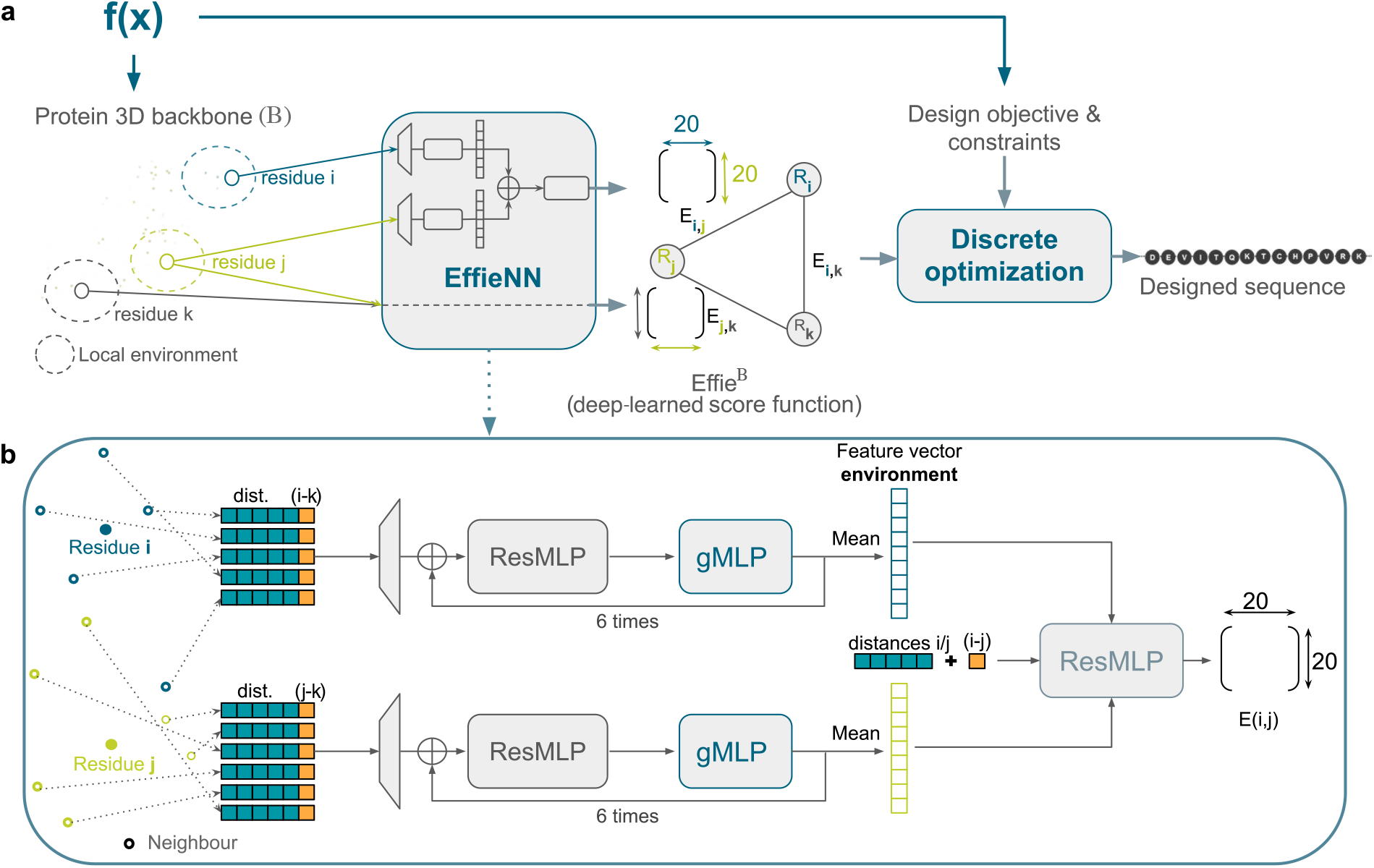
Overview of the generative neuro-symbolic AI protein design tool, EffieDes. **a** The coordinates of backbone atoms for each pair of residues (*i, j*) together with their neighbors in the input 3D protein structure *B*, are fed to the EffieNN neural network that predicts the pairwise interaction scores *E* (*s*_*i*_, *s* _*j*_ |*B*). The sum of all these pairwise terms defines a Potts model, Effie^*B*^, which can be further constrained with specific design objectives before optimization to design a protein sequence. **b** The neural network architecture of EffieNN is composed of several blocks, each comprising a multi-layer perceptron with residual connections (ResMLP) and a gated-MLP (gMLP). These blocks are applied six times to process a target residue and its neighbors, generating an environment embedding. A final ResMLP then outputs a 20×20 score matrix. The same neural network is used for all pairs of residues less than 15 Å away from each other.

The choice of Potts models provides a suitable inductive bias, as they have been shown to offer better representations of protein sequences than deep neural architectures such as variational autoencoders [35] or MSA-transformers [36].

Potts models offer both broad applicability and efficient integration within modular protein design workflows. They are widely used to represent coevolutionary couplings in protein sequence modeling [34, 42], to capture functional sequence fitness estimation derived from deep mutational scanning [43], and are also used to represent the score/energy functions over rotamers [44]in physics-based design tools, such as Rosetta or Osprey [45– 47]. Thanks to this shared mathematical form, they can all be easily combined with corresponding Effie^*B*^. Finally, being the description of the joint probability *P*(*s*_1_, …, *s*_*n*_ |*B*), Effie^*B*^ can be directly optimized, without the side-effect of low-temperature sampling, while the addition of constraints to the model enables retraining-free conditioning. Progress in automated reasoning over Potts models has given access to a variety of useful global constraints, enabling fine control over sequence composition [48–52], and complex design objectives [53].

Formally, the total score is decomposed into a sum of pairwise terms *E*_*i,j*_, representing interactions between residue pairs (*i, j*):

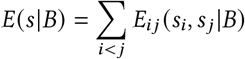

Each term *E*_*i,j*_ is predicted by EffieNN’s neural network as a 20×20 matrix representing local interaction scores for all possible choices of the corresponding pair of amino acids. The total score *E* (*s*|*B*) is trained to capture the probability of a sequence *s* given a backbone *B*, so that its optimization leads to maximally probable sequences for the input backbone structure. Furthermore, the positional identities required by structural symmetries or multi-state objectives [41] can be natively enforced within EffieDes through equality constraints.

Estimation of Potts models [54–57] is usually performed with Besag’s pseudo-loglikelihood (PLL) [58]. However, this loss function is known to struggle with highly unfavorable score terms [59], which are critical to capture strongly disfavored interactions. Therefore, we adopt the recently introduced E-PLL [60], which was specifically created to overcome this limitation. Training EffieNN with the E-PLL results in significant performance improvements over standard PLL (Figure 2).

**Figure 2:**
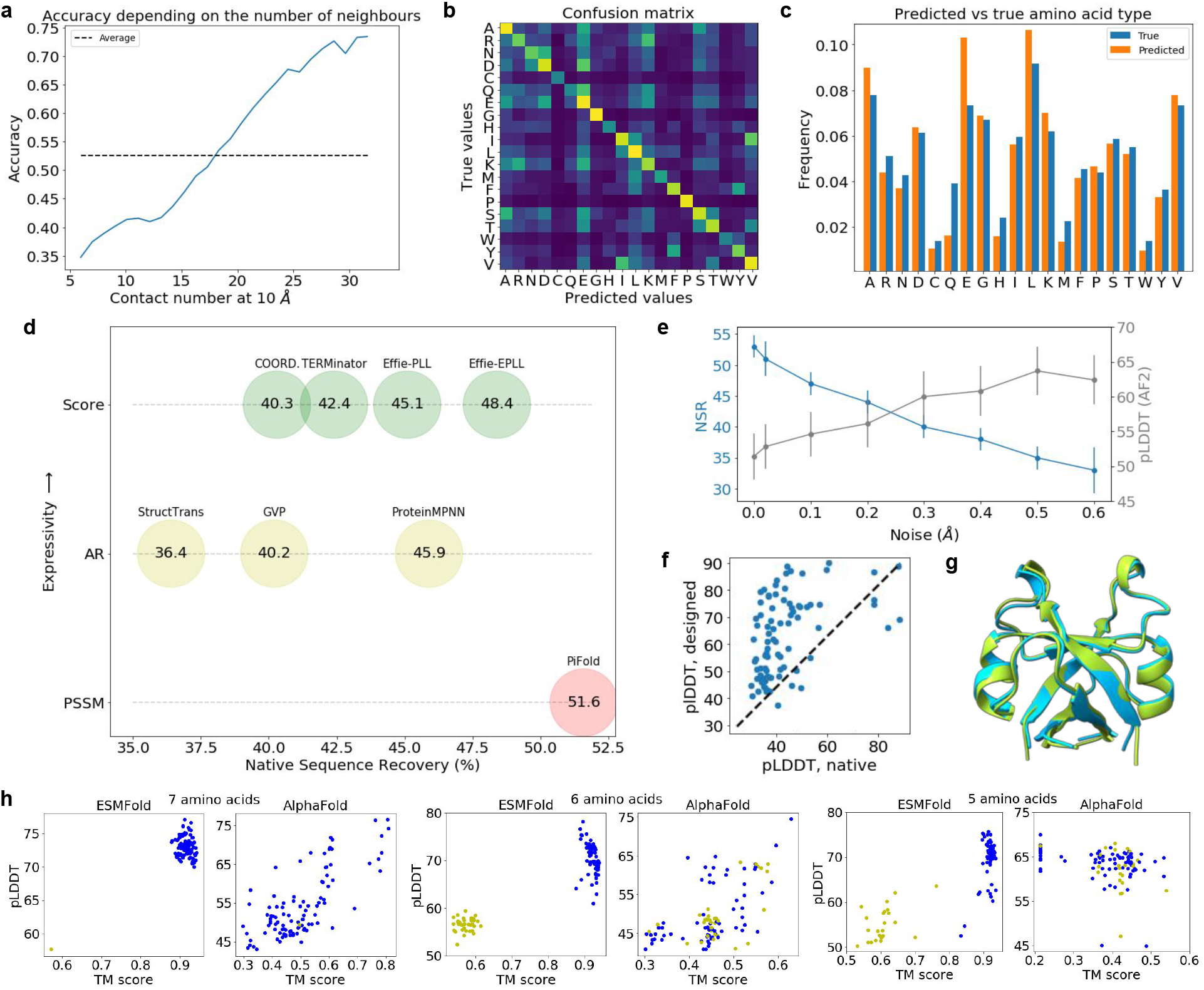
EffieDes predicts high-quality sequences from an input structure. For wide comparability, results in **a** to **d** were produced using only Ingraham’s single-chain dataset for EffieNN training. The remaining results were obtained using ProteinMPNN’s multi-chain dataset for training. **a** Prediction accuracy versus numbers of neighbors within 10 Å: buried residues with many neighbors are recovered more reliably than surface residues with few neighbors. **b** Confusion matrix: the highest confusions are observed for similar amino acids such as leucine (L) and isoleucine (I) or charged amino acids such as aspartic (D) and glutamic (E) acids. The lowest confusions are observed on amino acids with specific properties such as glycine (G), proline (P), or cysteine (C). **c** Histograms of predicted vs native amino acid types show very similar distributions. **d** Median NSR on the single-chain dataset design methods using deep learning (DL) methods. The compared methods either optimize a Potts model score (Score), decode the sequence auto-regressively (AR), or predict the most probable residue at each position (PSSM). Effie-PLL and Effie-EPLL refer to versions of EffieNN trained respectively using Besag’s pseudo-loglikelihood [58] or the E-PLL [60]. For consistency with other results [18], ProteinMPNN’s NSR ignores residues with missing backbone coordinates instead of using their native identity as originally reported [15]. CARBonAra, which reported a 51.5% NSR on monomers, is not included in Figure 2 because its training and test sets differ. **e** Training with perturbed coordinates decreases NSR but increases pLDDT: the error bars represent the standard error on a test set of 93 multi-chain proteins. **f** AlphaFold2 confidence, measured with pLDDT, in structure prediction of designed vs native sequences. Each dot represents a protein. **g** Target double *Ψ* -*β* barrel (DPBB) fold (green) aligned with the predicted fold (blue). **h** Forward folding metrics on all DPBB designs with 5, 6, or 7 different amino acids allowed, computed with ESMFold or AlphaFold. Blue dots are high-scoring sequences with ESMFold.

EffieNN is trained on structure-sequence pairs (*B, s*) derived from experimentally resolved protein structures. As local structural context largely determines amino acid identity, the model first extracts residue environments from the input backbone *B*, and then predicts interaction scores *E*_*ij*_ for each residue pair. The architecture of EffieNN, described in the Computational and Experimental Section and illustrated in Figure 1, combines residual multi-layer perceptrons (ResMLP) with gated MLPs [61], an efficient lean variant of Transformers.

We used two datasets for training. We first used the dataset proposed by Ingraham, Garg, Barzilay, and Jaakkola [62], which contains only single-chain structures and has been widely used in DL-based inverse folding methods, enabling fair comparisons. We then trained EffieNN using instead the multi-chain dataset of ProteinMPNN [15], restricted to soluble proteins. The ability of proteins to accommodate mutations through local backbone flexibility is accounted for by adding independent Gaussian noise to atomic coordinates during training.

### 2.2 EffieDes combines a deep-learned score function with automated reasoning for steerable protein design

Our generative neuro-symbolic AI framework EffieDes integrates Effie with the automated reasoning prover toulbar2 [39]. By framing protein design as a constrained optimization problem, EffieDes identifies sequences that maximize the predicted backbone fitness while strictly adhering to specific design objectives.

Although such constrained symbolic optimization enables high design expressivity, it also raises substantial computational challenges, as the resulting design problem is NP-hard [21]. To address this, EffieDes relies on different toulbar2 optimization algorithms that balance expressivity and scalability. For maximum expressivity, computationally-demanding exact algorithms provide optimal sequences satisfying complex additional constraints [63–67]. For large problems, the recent efficient polynomial-time LR-BCD algorithm [68] is preferred. LR-BCD solves a low-rank convex relaxation of the discrete optimization problem, an approach that can offer finite-time guarantees [69] on the expected optimized score, preferable to the asymptotic guarantees of stochastic optimization methods such as simulated annealing [45, 57]. Using data from Ingraham, Garg, Barzilay, and Jaakkola [62], we showed that LR-BCD matches the quality of exact methods while being significantly more scalable for practical applications.

Furthermore, for positive and negative multi-state design [41], we augmented EffieDes with a dedicated bi-objective optimization algorithm [53]. This extension enables the systematic exploration of Pareto-optimal trade-offs between competing design targets, allowing for a rigorous balance between desired structural affinity and necessary selectivity (see Supporting Information). It is also worth remembering that, as a Potts model, Effie should be directly usable in physics-based design platforms, such as Rosetta or Osprey [45–47], where it can be easily combined with physics-based score functions and fitness terms.

### 2.3 EffieDes predicts high-quality sequences from an input structure

Before applying EffieDes to full sequence design, we first assessed its ability to predict a single residue given all other amino acid identities within a given protein backbone *B* [56, 70]. This “masked residue prediction” task confirms the model’s grasp of local structural consistency. As expected (Figure 2a), buried constrained residues are recovered more frequently (73%) than surface-exposed residues (35%). The confusion matrix (Figure 2b) shows that substitutions predominantly occur between residues with similar physico-chemical properties. Conversely, amino acids with specific properties, such as proline, glycine, and cysteine, are predicted with high reliability. Because the predicted amino acid frequencies closely mirror natural sequences (Figure 2c), this indicates that EffieNN has successfully captured the intrinsic properties of natural proteins.

We then benchmarked EffieDes against other design approaches on full redesign problems, using the native sequence recovery (NSR) rate, the most common metric for assessing inverse folding performance. NSR measures identity between the designed and native sequences. We first compared EffieDes to methods that also exploit Potts models over rotamers or amino acids. On the same set of small proteins [62], EffieDes significantly outperforms Rosetta [45], achieving an NSR of 33.0% compared to 17.9% (results from Ingraham, Garg, Barzilay, and Jaakkola [62]). This result is particularly striking, given that Rosetta uses a detailed representation of side-chain atoms, while Effie relies on a learned implicit representation instead. As shown in Figure 2d, EffieDes also outperforms two approaches based on deep-learned Potts models [57]: COORDinator (based only on the structure coordinates, as Effie), and TERMinator [57], that additionally uses computationally expensive – 4 minutes per residue – Tertiary Repeating Motifs (TERMs). For these full redesigns of the single-chain test set, we ran EffieDes using the LR-BCD optimization algorithm.

Shifting to state-of-the-art auto-regressive methods — such as the Structured Transformer [62], GVP [14], and ProteinMPNN [15] — EffieDes demonstrates superior performance in terms of NSR. The theoretical capacity of auto-regressive models to learn high-order interactions [71] appears to be largely secondary in practice to the robust symbolic optimization employed by EffieDes. By directly sampling the joint distribution, EffieDes avoids the “greedy” pitfalls of sequential decoding. Moreover, unlike non-auto-regressive PSSM-based models such as PiFold [18], EffieDes provides the necessary framework for enforcing complex, interdependent constraints, making it a more versatile tool for advanced protein design.

We further validated EffieDes by assessing the quality of structure predictions of its designs using AlphaFold [72] in its single-sequence mode, thereby avoiding the confounding effect of multiple-sequence alignment (MSA) depth variability. As shown in Figure 2f, EffieDes-designed sequences generally substantially surpass native sequences in pLDDT scores. This shows that EffieDes encodes structural information more effectively than naturally occurring sequences, which must typically balance multiple evolutionary pressures. Additionally, we found that introducing Gaussian noise to atomic coordinates during training (up to 0.5 Å standard deviation) slightly trades off sequence recovery for significant gains in AlphaFold confidence (Figure 2e), a trend previously observed in other state-of-the-art models [15].

### 2.4 EffieDes can condition sequence design without retraining

Functional protein design frequently requires to enforce complex sequence constraints, ranging from motif preservation and reduced amino acid alphabets to strict structural symmetries and multi-state requirements. Enforcing these constraints typically follows one of three paths. First, one can retrain the model on specialized datasets — a human and computing-intensive process which may even be infeasible for objectives for which no associated data exists. Second, auto-regressive models can modulate local conditional probabilities *P* (*s*_*i*_|*s*_*<i*_, *B*) to favor specific residues or combine the distributions of symmetrical/identical residues [15]. This “greedy” adjustment causes the sampling to diverge from the foundational chain rule, with unpredictable consequences for sequence validity. Third, the neuro-symbolic approach of EffieDes incorporates these constraints directly into the global optimization of the learned Potts model Effie. By adding global constraints — including amino acid composition, structural symmetries, and multi-state objectives – to a learned Potts model, EffieDes enables “zero-shot” sampling in regions of the fitness landscape for which no training data exists.

A striking example of this appears in the work of Yagi, Padhi, Vucinic, et al. [73]. To show that the double-Ψ *β*-barrel (DPBB), a key structural component of RNA polymerase, could have originated from an early translation system and genetic code, they redesigned a DPBB sequence with a bounded number (7) of distinct amino acid types. We build on this work to illustrate the benefits of EffieDes’s conditioning capabilities. We used EffieDes to redesign the DPBB fold while imposing even stricter constraints, limiting the number of amino acid types to *k* = 7, 6, or 5. This is a particularly challenging problem, as it requires generalization beyond the training distribution, no known globular protein being composed of such a restricted amino acid set. Given that the final set of amino acid types to use is unknown, it is unclear how an auto-regressive model could tackle such a design problem.

Since the DPBB forms a symmetric dimer, the sequence was optimized under a symmetry constraint of order 2 and a dedicated *N* -value global constraint [50, 74], limiting the number of different amino acid types used overall. We designed 100 sequences for each value of *k*. For *k* = 7, these sequences use 3 different subsets of amino acids, including the experimentally validated {D, A, G, R, V, E, K} set [73]. They share a consistent composition, systematically including the amino acids D, A, G, R, and V. The structure of each design was then predicted using AlphaFold [72] in single-sequence mode and ESMFold [9] and evaluated using the confidence scores (pLDDT) of each prediction tool and alignment to the target fold score (TM).

Despite the out-of-distribution input sequences, ESMFold accurately predicted the *k* = 7 folds, with TM-scores close to 1.0 for all sequences except one (Figure 2g-h). AlphaFold produced more scattered metrics, including high-quality sequences with TM *>* 0.75 and pLDDT *>* 75. AlphaFold is not specialized for single-sequence structure prediction, which may explain its poorer performance here. For both models, there is a clear correlation between pLDDT and TM scores, except for AlphaFold on the most challenging case of *k* = 5. Although reducing the amino acid repertoire expectedly complicates the design landscape, ESMFold continues to discern high-quality, foldable sequences even for *k* = 5 (Figure 2h).

By leveraging automated reasoning to optimize Potts models under complex constraints, EffieDes can steer sampling into fitness regions inaccessible to auto-regressive methods. This bypasses the need for retraining — a process that is unfeasible here precisely because no training data exists for the specific design objective. This makes EffieDes a powerful tool for exploring a ’dark’ sequence space that remains out of reach of purely datadependent architectures.

### 2.5 EffieDes can design symmetrical multi-component assemblies

Protein assemblies are crucial for many biological processes [75, 76], and the ability to design specific protein-protein interactions opens up new opportunities for progress in a wide range of fields, from fundamental science to health and biotechnology [77–81]. A significant challenge in this area lies in precisely controlling the interactions between protein components to achieve the desired assembly. This challenge becomes even more critical when designing symmetrical assemblies composed of sequence-distinct proteins sharing a common fold.

To evaluate the ability of EffieDes to handle such complex settings, we applied it to the redesign of bacterial microcompartment shell proteins (BMC-H), which natively form homo-hexamers [82–85] (Figure S6, Supporting Information), into hetero-hexamers composed of two distinct subunits (*e.g*. **A** and **B**, Figure 3a). Such a redesign is attractive from a biotechnological perspective, as hetero-assemblies could expand the functional versatility of BMC scaffolds by potentially enabling asymmetric catalytic arrangements, tunable permeability, or modular surface display [86–98]. However, achieving such controlled hetero-assemblies requires the simultaneous enforcement of symmetry constraints and a multi-state design strategy [99, 100], in which the heteromeric state (AB, positive) is favored while the homomeric states (AA and BB, negative) are avoided (Figure 3a).

**Figure 3:**
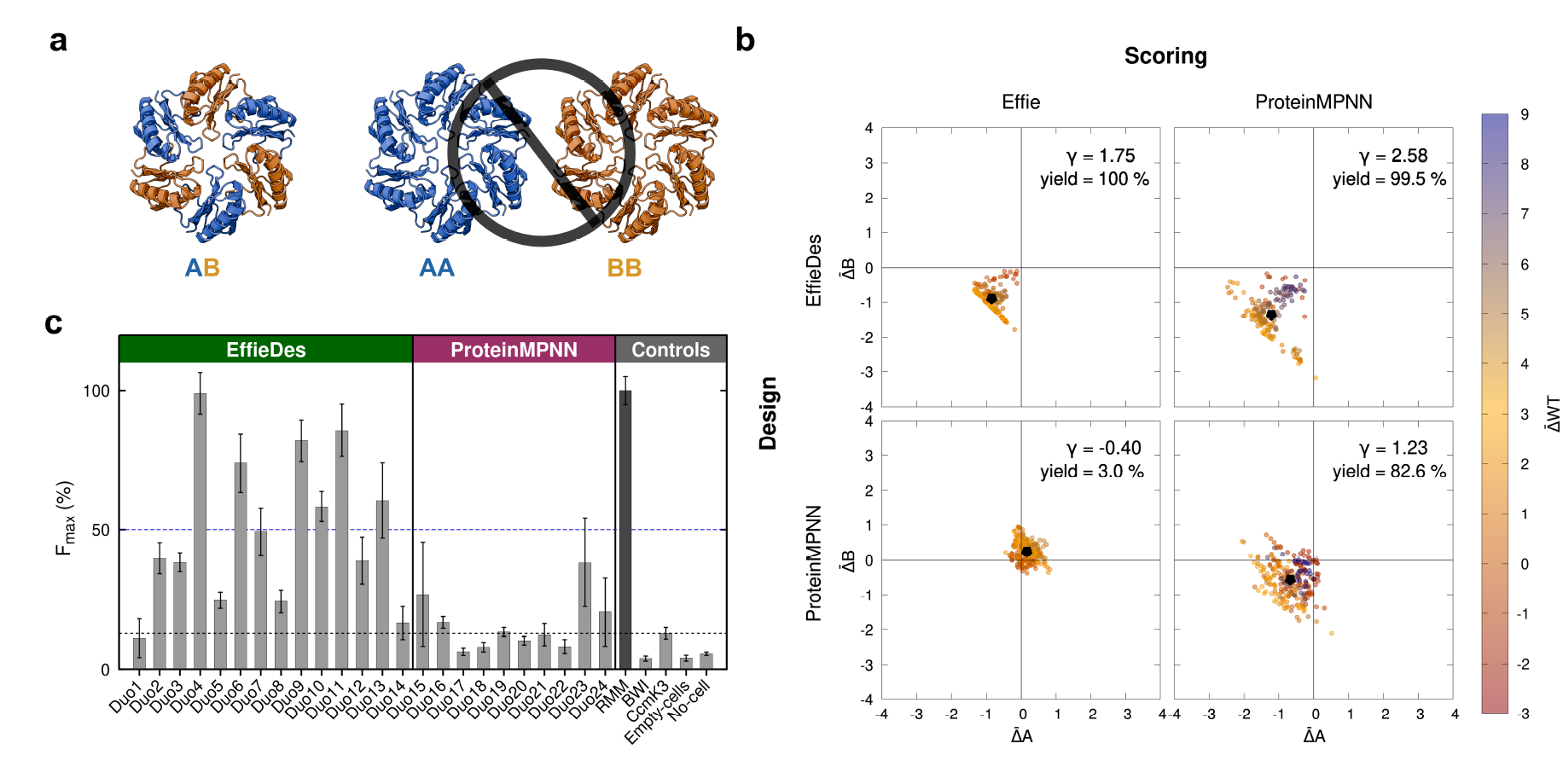
Design of protein interfaces in self-assembling heteromeric BMC-H proteins. **a** Representation of the negative design strategy for BMC-H proteins showing the heterohexameric positive state (AB) and the two homohexameric negative states (AA and BB). **b** Comparison of the pre-minimization score differences between positive and negative hexameric states for mutant BMC-Hs designed with EffieDes (top row) or ProteinMPNN (bottom row), using the small designable region. 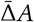 and 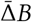 *B* are represented on the *X* and *Y* axes and each dot represents a design. The color scale indicates the 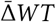 scores, with blue corresponding to heterohexamers (AB) that are likely weakly stable. Regardless of the scoring approach used, EffieDes designs exhibit better 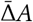 and 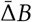, as highlighted by the averaged 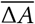 and 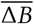 (black dots) and their γ values. **c** Experimental screening of the associations between designed pairs of BMC-H monomers A and B using the tripartite GFP technology. Maximal fluorescence (*F*_*max*_) values are given as a percentage of the WT RMM value (dark gray). The dashed black line highlights maximal signals emitted by negative controls: non-associating pairs (buckwheat trypsin inhibitor or BWI), aggregating proteins (CcmK3 from *Synechocystis sp. PCC 6803*), or cell auto-fluorescence (Empty: cells transformed with a pET26b empty vector).

A 3D symmetrical hexamer design template was built from the crystal structure of the BMC-H shell protein RMM (PDB ID=5L38) [82], and two designable interfacial regions (small and large) were defined (see Computa-tional and Experimental Section and Figure S7, Supporting Information). Both EffieDes and ProteinMPNN [15] were tasked with generating sequence pairs A and B that satisfy the negative design objective of selective heteromeric interaction (see Computational and Experimental Section and Supporting Information). A total of 377 pairs (AB) of sequences were designed using EffieDes and 495 using ProteinMPNN. To quantitatively evaluate how effectively each method favored the hetero-hexamer over the homo-hexamers, all sequences were cross-scored by both Effie and ProteinMPNN, and the normalized differences in score of the positive and negative states were computed (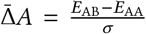 and 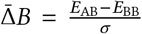, where σ represents the standard deviation of all Δ*A* and Δ*B* values; see Computational and Experimental Section). Similarly, the stability of the heterohexamer AB relative to the wild type (WT) homomeric template was assessed by computing 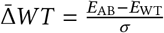.

All the pairs of sequences designed by EffieDes show a better score for the positive state AB than for the negative states AA or BB as highlighted by the negative 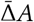 and 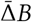 Effie scores. In contrast, ProteinMPNN is unable to completely account for the negative design objective, as only *∼*85% of its designs show negative 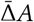 and 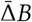 ProteinMPNN scores (Figure 3b and Figures S8-11, Supporting Information). This trend is further captured by the composite metric 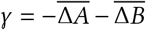 (with 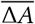 and 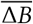 the averaged values, see Computational and Experimental Section), which quantifies progress along the intended optimization direction. Indeed, γ_*EffieDes*_ *>* γ_*ProtMPNN*_ regardless of the scoring method used for evaluation (Table S2, Supporting Information). Strikingly, ProteinMPNN gives better 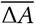 and 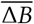 to EffieDes designs than to its own designs (Figure S11, Supporting Information). This underlines the incapacity of auto-regressive models to sample high-quality sequences when strong conditioning requires the sampling of conditional distributions that drift far away from those specified in the fundamental chain rule.

As a decomposable function, Effie also provides valuable insights into how mutations affect monomer interactions by separating intra- and inter-chain scores. The Δ*A*_inter_ and Δ*B*_inter_ score differences (Figures S12 and 13, Supporting Information) further highlight the limitations of ProteinMPNN in solving this negative design problem, as its γ_inter_ is strictly negative (Table S3, Supporting Information).

The stability of all designed hexamers was investigated by constructing and minimizing the structures of all positive and negative states using the Rosetta beta_nov16 score function [101] (Figures S9-15, Supporting Information). Candidates from either approach were then selected for experimental testing based on their favorable Effie or ProteinMPNN score differences Δ*A*, Δ*B*, and Δ*WT* post-minimization, as well as low root-mean square deviation (RMSD) of the positive state AB backbone to the WT hexamer (see Computational and Experimental Section, Supplementary Discussion and Tables S4-7, Supporting Information). The tripartite-green fluorescent protein (tGFP) technology [102, 103] was used to detect AB interactions in vivo upon co-expression in *Escherichia coli*. This method relies on a GFP reporter split into three non-fluorescent portions, two of them being attached respectively to A or B. The reconstitution of the complete GFP in an environment containing the third missing GFP component, with concomitant fluorescence emission, requires the association of the two protein partners (A and B) (Figure S17, Supporting Information).

The experimental success rate of EffieDes was markedly higher than that of ProteinMPNN, with 86% (12/14) of designs exhibiting fluorescence levels well above negative controls compared to just 20% (2/10) for Protein-MPNN (Figure 3c). Furthermore, while nearly half of the EffieDes designs (6/14; 43%) exceed 50% of the WT RMM positive control’s fluorescence, this level of performance was not reached by any ProteinMPNN-designed sequence (0/10). Analysis of protein contents by SDS-PAGE corroborates the superior efficiency of EffieDes, as ProteinMPNN designs more frequently result in aggregation of a fraction of the expressed mutant proteins (see Supplementary Discussion and Figure S16, Supporting Information). Among all candidates, the “Duo4” design from EffieDes shows the highest expression and interaction between AB protein partners. This is supported by fluorescence measurements after comparison of protein expression levels (Figure 3c) and by co-purification experiments (see Supplementary Discussion and Figures S17-18, Supporting Information). Moreover, the formation of Duo4 hexamers was confirmed by size-exclusion chromatography (SEC) experiments, as well as their stability (see Supplementary Discussion and Figure S19, Supporting Information).

EffieDes thus successfully designed new BMC-H proteins that self-assemble in vivo. Overall, in silico and experimental results indicate that EffieDes can better address complex negative design problems involving symmetric objects than the auto-regressive ProteinMPNN method for new associations of pairs of mutant BMC-H proteins.

### 2.6 EffieDes can design nanobodies using de novo backbones

Nanobodies, also known as VHHs (single-domain antibody fragments derived from the variable domain of heavychain-only antibodies found in camelids), are increasingly used in biotechnology and therapeutics due to their ease of production, high stability, and ability to access cryptic epitopes. Engineering nanobodies using computational design methods can offer a fast and scalable alternative to conventional discovery approaches, such as animal immunization and the screening of immune or synthetic libraries. However, the structure-based design of nanobodies remains challenging, as it requires an accurate modeling of their complementarity-determining regions (CDRs), highly variable and functionally critical loops that govern antigen-binding specificity and affinity [104]. While generative AI approaches, such as RFdiffusion [13, 105], can, in principle, de novo generate CDR backbones tailored to an antigen surface, the challenge remains to identify sequences that effectively encode these pre-organized geometries while simultaneously optimizing the interfacial interactions for affinity and target specificity. This task is particularly demanding for highly flexible CDR loops, where small sequence changes can profoundly reshape the conformational and energetic landscape of the binding interface. Here, we evaluated whether the global optimization capabilities of EffieDes could design functional sequences for novel CDR backbones targeting an immune-evasive virus.

We centered our study on MR17, a synthetic nanobody (sybody) originally developed against the ancestral Wuhan strain of SARS-CoV-2, which binds to the receptor-binding domain (RBD) of the spike protein and blocks its interaction with the human angiotensin-converting enzyme 2 (ACE2) receptor [106]. It retains binding to the Delta variant RBD, but fails to interact with the Omicron XBB.1.16 variant, which carries 11 mutations at the MR17 binding interface compared to the Wuhan strain (Figures S27-28, Supporting Information). As MR17 targets a functionally critical epitope, it provides a relevant structural template to assess whether EffieDes-based sequence design on de novo CDR backbones can rapidly retarget a nanobody toward an emerging immune-evasive variant.

To achieve this, we generated novel CDR backbones for MR17 [106] in complex with the SARS-CoV-2 XBB.1.16 RBD by partial diffusion using RFdiffusion [13] and subsequently designed sequences by either ProteinMPNN sampling or EffieDes optimization, using global diversity constraints [49, 52] to ensure a systematic exploration of the highest-fitness regions of the landscape. A total of 5, 600 sequences were sampled with ProteinMPNN and independently, 90 optimized with EffieDes. These designs were evaluated primarily based on predicted ΔG binding affinities computed with Rosetta’s InterfaceAnalyzer and the beta_nov16 score function [101, 107] (Table S10, Supporting Information), and with MM/GBSA and MM/PBSA binding energy estimates [108–112] from molecular dynamics (MD) simulations (see Supplementary Methods, Supporting Information). Additional structural criteria, such as interfacial hydrogen bond propensity and interface size, were also considered throughout the MD trajectories (Tables S11-12, Supporting Information). Nine nanobody candidates – eight from Protein-MPNN and one from EffieDes– were selected for experimental testing (Figure S30 and Table S13, Supporting Information).

Of the nine designs tested, only the EffieDes-derived nanobody (NbRM-E1, Figure 4a) exhibited binding to the SARS-CoV-2 XBB.1.16 RBD in biolayer interferometry (BLI) experiments (Figure S21, Supporting Information). Notably, NbRM-E1 binds to its target with a higher affinity than MR17 to the Wuhan RBD [106]: K_D MR17_ =83.7 nM with k_off MR17_ = 9.7 10^−3^ s^−1^ *vs* K_D NbRM-E1_ = 64 nM with k_off NbRM-E1_ = 4.74 10^−3^s^−1^ (Figure S24, Supporting Information). Furthermore, while MR17 interacts with the RBD of the SARS-CoV-2 Delta variant, NbRM-E1 selectively binds to the XBB.1.16 RBD, with no detectable interaction with the Delta RBD (Figures S25-26, Supporting Information).

**Figure 4:**
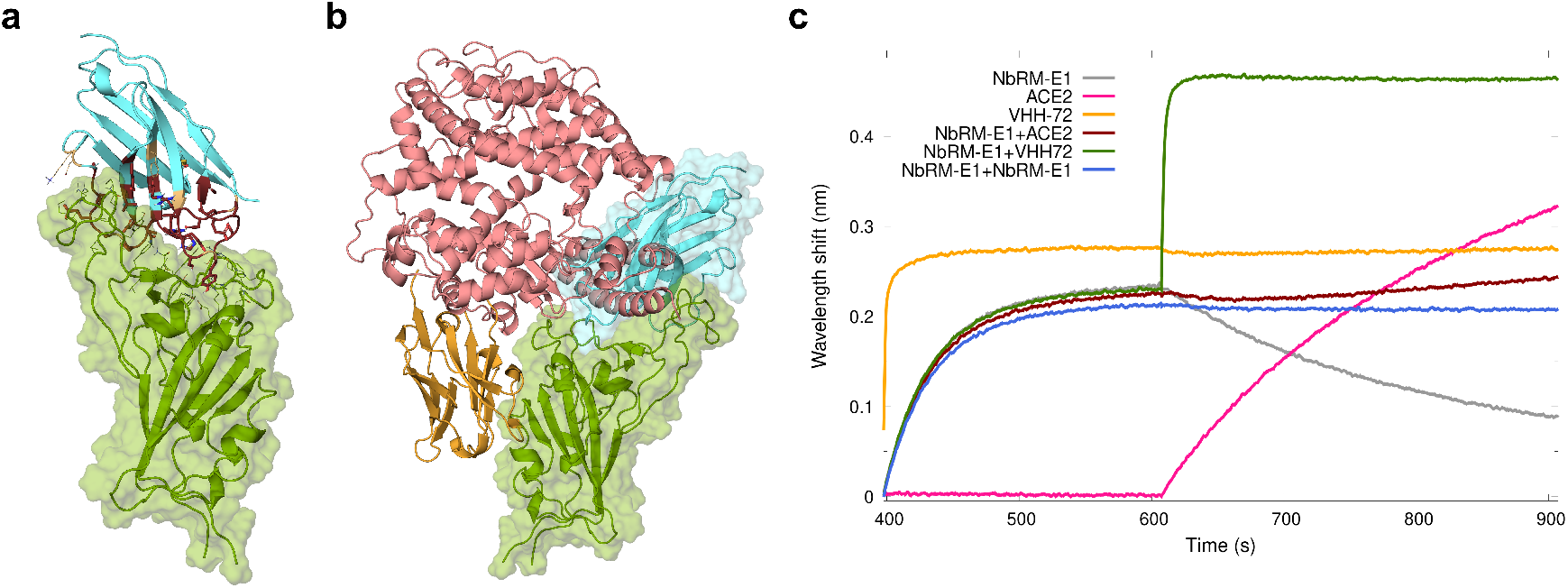
Design of nanobodies binding the RBD of SARS-CoV-2 XBB.1.16. **a** 3D structural model of nanobody NbRM-E1 (cyan) in complex with the SARS-CoV-2 XBB.1.16 RBD (green). Interfacial residues are represented as lines with, on NbRM-E1, the 7 native designable residues colored in orange and the 29 residues distinguishing NbRM-E1 from MR17 shown as dark brown sticks. **b** 3D model showing the binding sites of each protein partner (NbRM-E1: cyan, ACE2: pink, and VHH-72: orange) on XBB.1.16 RBD (green), generated by aligning the backbone of the XBB.1.16 RBD with that of the RBD in complex with ACE2 (XBB.1.5, PDB ID: 8VKP) [113] or with VHH-72 (SARS-CoV-1, PDB ID: 6WAQ) [114]. **c** BLI competition assays showing the inhibition of ACE2 binding when NbRM-E1 is interacting with XBB.1.16 RBD (dark brown) and the binding of VHH-72 nanobody to the complex NbRM-E1/RBD (green), which supports the predicted binding site of NbRM-E1.

BLI-based competition assays (see Computational and Experimental Section) show that the NbRM-E1 nanobody effectively blocks the interaction between the XBB.1.16 RBD and its ACE2 receptor. Furthermore, NbRM-E1 allows for the simultaneous binding of VHH-72—a broad neutralizing nanobody targeting a conserved, non-overlapping epitope [114, 115] (Figure 4b-c). This dual observation—ACE2 blockade coupled with a lack of competition with VHH-72—provides a robust functional mapping that aligns with the binding interface proposed in our structural model (Figure 4a-b).

EffieDes has thus proven its ability to design a functional sequence on a de novo CDR backbone that enables binding to the SARS-CoV-2 XBB.1.16 RBD, thereby successfully retargeting it toward an immune-evasive viral variant. The combination of backbone generation and EffieDes sequence design thus provides a rapid and effective strategy for engineering nanobodies in response to emerging viral threats.

## 3 Discussion

The rapid ascent of deep learning for inverse folding has largely been driven by the efficiency of auto-regressive architectures. However, our findings suggest that the reliance on sequential, token-by-token decoding — effectively a “Next-Token Prediction” approach — imposes a structural ceiling on design complexity. In inverse folding, where long-range side-chain coordination is paramount, the inability of these models to “think ahead” leads to an accumulation of local errors. By replacing sequential sampling with the global optimization of a backbone-dependent Potts model, EffieDes restores the ability to account for the intricate, long-range inter-residue interactions inherent in protein folding. This shift from a “greedy” local walk to a global *Maximum a Posteriori* inference allows for the exploration of fitness landscapes that are mathematically unreachable by current auto-regressive standards.

The most significant advantage of the neuro-symbolic framework is its capacity for “hard” conditioning without the need for specialized retraining. Our results on the BMC-H hexameric platform provide a clear demonstration of this: while ProteinMPNN struggled to balance the competing requirements of symmetry and negative design, EffieDes identified viable sequences by treating these requirements as formal logical invariants. The fact that ProteinMPNN, during in silico cross-evaluation, favored sequences designed by EffieDes over its own suggests that the failure of auto-regressive models in these settings is not a lack of ‘knowledge’ within the weights, but a failure of the sampling process itself under strong conditioning. This “sampling breakdown” occurs because, as constraints are added, the effective search space narrows to a region that the sequential chain rule can no longer accurately approximate.

Furthermore, the successful design of the NbRM-E1 nanobody against an immune-evasive SARS-CoV-2 variant highlights a second critical strength: data efficiency. By utilizing the inductive bias of a Potts model – which naturally captures the pairwise nature of physical interactions – EffieDes demonstrates an ability to generalize to de novo CDR loop structures that exist outside the training distribution. This suggests that neuro-symbolic architectures may be less prone to the ’memorization’ pitfalls of pure deep learning, offering a more robust path for designing proteins for which evolutionary data is sparse or non-existent. The nanomolar affinity and ACE2-blocking capability of NbRM-E1 confirm that this theoretical advantage translates directly into functional, high-performance molecular tools.

To further expand the reach of EffieDes, several architectural evolutions can be envisioned. While current inverse folding tools are beginning to incorporate non-protein ligands [16, 22], their authors acknowledged the limitations of pure DL methods, particularly in low-data regimes, suggesting their hybridization with physics-based methods. Effie score functions would be a fitting candidate as they can already be combined with physics-based pairwise score functions for ligands, directly or using a new all-atom version of EffieNN. Finally, while the current pairwise Potts representation is highly effective, extending EffieNN to capture many-body interactions could further refine the resolution of the fitness landscape, paving the way for the design of increasingly sophisticated molecular machines.

## 4 Computational and Experimental Section

### Protein structure representation and dataset

The input protein backbone *B* is defined by the 3D coordinates of *N, C*_*α*_, *C*, and *O* atoms for *n* residue, such that *B* ∈ *R*^*n*×4×3^. To enforce the SE(3) (global rotation and translation) invariance of protein structures [116] EffieNN encodes backbone geometry using invariant pairwise features between residue pairs (*i, j*):

- Interatomic distances: for each residue pair (*i, j*), all pairwise distances between the backbone atoms (*N, C*_*α*_, *C, O*, and virtual *C*_*β*_) of residues *i* and *j* are computed [15]. The resulting 25 distances are encoded using 16 Gaussian radial basis functions with centers evenly spaced between 0 and 20 Å.
- Positional encoding: a 16-dimensional sinusoidal embedding is used to encode the number of residues between positions *i* and *j* in the sequence (| *j* − *i* |) [117].
- Chain identity: for multi-chain proteins, a Boolean feature indicates whether residues *i* and *j* belong to the same chain, to learn interfacial properties [118].

The final representation of the protein backbone is a matrix of pairwise features with shape (*n, n*, 25×16+ 16+1). Residues with missing atomic coordinates in experimentally resolved structures are not predicted and excluded from all metrics.

EffieNN was trained on two datasets extracted from the Protein Data Bank [20]. The first *single-chain* dataset, assembled by Ingraham, Garg, Barzilay, and Jaakkola [62], is composed of single-chain protein structures. Training, validation, and test sets (respectively, 17,000, 600, and 1,200 proteins) are split based on a sequence similarity cut-off of 40 % and CATH 4.2 classification to ensure that similar proteins are in the same set. This dataset has been widely used to train and test inverse folding architectures, allowing for fair comparison across methods. The second *multi-chain* dataset, used for ProteinMPNN training, is composed of protein assemblies clustered into training (23,358), validation (1,464), and testing (1,539) sets. Each assembly can be associated with multiple sequences and/or conformations, one of which is randomly sampled at each training epoch. This dataset overcomes a key limitation of the single-chain dataset, which can fragment biologically relevant complexes into isolated chains and overlook interfacial properties. Only soluble proteins shorter than 1, 000 amino acids were retained.

### Neural architecture and training

EffieNN comprises two main components, as illustrated in Figure 1. The first component extracts information from the local environment of each residue, defined by its 128 nearest neighbors. For small proteins smaller than 128 amino acids, zero-padding is applied. The environment is processed using blocks composed of a gated Multi-Layer Perceptron (gMLP) [61] and an MLP with residual connections [119] (ResMLP). Inspired by Graph Neural Network architectures, the same gMLP+ResMLP bloc is repeated 6 times, with the output of the previous iteration added to the input. The final output is averaged over the neighbors to produce an embedding of the target residue’s environment. The second component, a single ResMLP, predicts pairwise score matrices *E*_*i,j*_. For all pairs of neighboring residues (the *k* = 128 nearest neighbors), the concatenated environment vectors of each residue in the pair, along with the distance and positional features, are fed to this ResMLP to produce a 20×20 matrix. The set of all such pairwise matrices defines the generated Effie^*B*^ score function, conditioned on the input backbone *B*. An ablation study and detailed hyperparameters are available in the Supporting Information.

EffieNN training is self-supervised to maximize the probability of the sequences observed in the training set given their associated backbone. The exact loglikelihood criteria being intractable for Potts models, it is usual to use Besag’s pseudo-loglikelihood (PLL [58]). We rely instead on the E-PLL [60], which avoids the known incapacity of the PLL to estimate high-energy terms [59] by randomly masking a fraction of pairwise terms during training. To accommodate variable protein sizes, the E-PLL eliminates 10 % of the pairwise terms connecting a residue to its neighbors during training. A L1 regularization of 10^−4^ is applied to the predicted score matrix to encourage sparsity and simplify discrete optimization. Following preliminary training that showed constant zero functions estimation for pairs of distant vertices, and consistently with physics-based score functions, no function is predicted for pairs of residues separated by more than 15 Å.

EffieNN was trained using the Adam optimizer, with a weight decay of 10^−3^ and an initial learning rate of 5.10^−4^. The learning rate is divided by 10 when the validation loss decreases (with patience 0) until it reaches 10^−8^. Each protein in the training set is treated as a batch of *n* residues (of varying size). This variation becomes important in the multi-chain dataset, with proteins having from 50 to 1,000 amino acids: the learning rate is set proportionally to the square root of the batch size [120].

### In silico evaluation of EffieDes

EffieNN is optimized by the automated reasoning prover toulbar2 [39]. Three algorithms are used: exact optimization with the Hybrid Best-First Search algorithm [121], low-rank convex relaxation optimization LR-BCD [68] for efficient solving, and a custom bi-objective solver developed for multi-state design. Exact optimization was used for benchmarking against Rosetta on small proteins and for the redesign of RNA-polymerase, using toulbar2’s Python interface, PyToulbar2 (version 0.0.0.2). For large proteins (*e.g*., NSR comparisons on the full single-chain dataset), optimization was performed using LR-BCD with 50 rounding runs, 5 iterations, and option −4 for the rank. The sequence with the lowest predicted score among the 50 produced was retained. LocalColabFold [122] (version 1.5.2) with 3 recycles was used to assess the quality of the designed proteins through forward folding.

### RNA polymerase redesign

The crystal structure of a previously designed double-*Ψβ*-barrel (DPBB), composed of seven amino acid types [73] (PDB ID: 7DXZ) was used as a starting point. The symmetry axis between the two DPBB monomers was determined using AnAnaS [123], and the backbone was idealized to enforce perfect symmetry with Rosetta. Each monomer consists of 42 amino acid residues. Four Effie score functions were predicted by 4 versions of EffieNN trained on the single-chain or multi-chain dataset, with or without added backbone noise of 0.2 Å during training and optimized exactly with toulbar2. To constrain the number *k* of amino acid types allowed in the design, an *N* -value constraint [74] was encoded as a generalized linear constraint [50] in toulbar2, with auxiliary Boolean variables indicating whether each amino acid type is used. Without any constraint, the optimal sequence uses 12 different types of amino acids among the 20 available. Constrained design was performed for *k* ∈ 5, 6, 7 amino acid types by enumerating all sequences within 10 units of the optimal score. The 100 best-scoring sequences (per *k* and per Effie model) were retained for downstream analysis; for *k* = 5, all valid sequences found (fewer than 100) were kept.

### Implementation of multi-state design approaches for the redesign of BMC-H proteins

We used EffieNN trained on the multi-chain protein dataset, with 0.14 Å std-dev Gaussian noise added to the atomic coordinates during training. EffieNN independently processed both the positive and negative states, and the resulting decomposable score function was projected on monomers based on the symmetrical properties of each state. The approximate Pareto front was produced for two designable regions of respectively 29 and 39 residues, as described in Supporting Information. This produced 450 (resp. 778) solutions for the small (resp. large) region (Figure S4, Supporting Information). Following sequence clustering, the sequence minimizing the absolute value of the difference in Effie scores between the two negative states, AA and BB, was selected in each cluster, producing 210 and 204 unique sequences for evaluation. To favor optimization of the positive state, only sequences for which |*λ*^−^| *< λ*^+^ were kept, with *λ*^−^ the weight applied on the negative states and *λ*^+^ the one applied on the positive state, resulting in a final count of 189 (resp. 188) sequences.

ProteinMPNN was used with its default parameters: `model v_48_020`, trained with 0.20 Å Gaussian noise, and a sampling temperature *T* = 0.1. The positive state (AB) was assigned a weight *λ*^+^ = 1. As with EffieDes, to give more importance to the positive state, weights *λ*^−^ ranging from −0.9 to −0.1 by increments of 0.1 were tested for the negative states (AA and BB), using the same weight for both states. Symmetrical chains in all states were tied using the `homooligomer` option of the `make_pos_neg_tied_positions_dict.py` script of ProteinMPNN. Combinations with a negative weight between −0.45 and −0.25 were determined to be the most suitable for our design objective and were explored with a finer resolution of 0.01 (see Supporting Information). For each weight combination, 10 designs were generated, for a total of 280 pairs of sequences for each designable region. Duplicate sequences were removed, leading to a total of 236 (resp. 259) unique designs.

### Construction of conformations and in silico assessments of the redesign of BMC-H proteins

After identification of the symmetry axis of the X-ray structure of the BMC-H shell protein MSM0272 forming the *Rhodococcus* and *Mycobacterium* microcompartment (RMM) from *Mycobacterium smegmatis* MC^2^ 155 (PDB ID=5L38) [82] using AnAnaS [124], Rosetta cyclic C6 symmetry definition [125] was used to build a symmetrical hexamer conformation, with a 60° rotation between each asymmetric subunit and a distance of 17.1 Å between the center of mass of the subunits and the center of the cyclic hexamer. This hexamer was then minimized using Rosetta `ref2015` Cartesian scoring function [126] with the same symmetry constraints. The resulting conformation is denoted as the wild-type (WT) template and was used for the design of heteromers AB (Figure 3a) and as a reference for design evaluation. A small (29 residues) and a larger (39 residues) designable region were defined at the interface between the subunits (Figure S7, Supporting Information). Mutant conformations of the three different states considered in our multi-state designs (one heterohexamer AB and two homohexamers AA and BB) were built by mapping the designed sequences on the WT template conformation using our in-house Custozyme+ tool, side chain packing all residues using Rosetta `beta_nov`16 scoring function [101]. These mutant conformations were then minimized using the Rosetta minimize executable with the multi-step L-BFGS algorithm (`lbfgs_armijo_nonmonotone`), a minimization tolerance of 0.001, and beta_nov16 scoring function. No symmetry constraints were applied, allowing for large conformational changes in the hexameric conformations (Figures S12-13, Supporting Information). Conformations pre- and post-minimization were scored using both Effie and ProteinMPNN and the scores differences between the positive (AB) and each negative (AA or BB) hexameric state Δ*A* = *E*_AB_− *E*_AA_ and Δ*B* = *E*_AB_− *E*_BB_ were computed, as well as those between the positive state and the wild-type homomeric RMM template Δ*WT* = *E*_AB_−*E*_WT_. To account for possible discrepancies in the score units from Effie and ProteinMPNN, score differences were normalized by the standard deviations over all

Δ*A* and Δ*B* values from Effie scores or from ProteinMPNN scores (σ_Effie_ = 171.134, σ_ProtMPNN_ = 64.1453). For a design scored with Effie, this defines 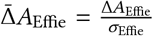. The averaged values 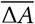 and 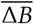 were calculated for each approach (*e.g*. for each design *i*, 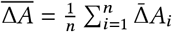, as well as the scalar product 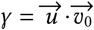 with 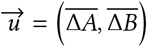 and 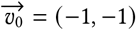. Similarly, the Effie inter-chain score differences were calculated for all designs, and normalized by the standard deviation on all Δ*A*_inter_ and Δ*B*_inter_ values (σ_inter_ = 148.985), as well as the averaged values and the scalar product γ_inter_.

### Production and characterization of the designed BMC-H proteins

Full-length DNA fragments coding for either Duo-GFP10 or Duo-GFP11 were synthesized by Twist Bioscience (sequences provided in Table S8, Supporting Information). Each pair of fragments was then Gibson-assembled within a NdeI/SalI-treated pET26b-based vector-1 (Kan^R^) that also coded for an His_6_-tagged GFP1-9 under an independent T7-transcription control. Gibson assembly reactions were carried out with the NEBuilder Hifi DNA Assembly Master Mix (E2621, NEB) following supplier instructions. Reactions were transformed in TOP10 cells and plasmids recovered by standard purification procedures [103]. Final vectors were verified by restriction and sequencing (Eurofins).

Protein expression studies were performed after the transformation of chemically-competent BL21(DE3) *E. coli* cells with the plasmids. A preculture of each strain in Luria-Bertani broth (LB, 100 µL) media supplemented with kanamycin (40 µg mL^−1^ final conc.: LBK media) was grown for 6 to 8 h at 37 ^°^C and 200 rpm orbital-shaking. Then, 2 µL were seeded into LBK (200 µL) supplemented with IPTG (10 µM), previously dispensed in a 96-well black plate with glass flat bottom (Greiner, reference 655892). Cultures were continued in a CLARIOstar Plus (BMG Labtech), at 37 ^°^C with continuous shaking. Optical density (at 600 nm) and GFP fluorescence (excitation wavelength at 470 ± 15 nm, emission at 515 ± 20 nm) were recorded approximately every 10 min, for 15 h. Data were fitted to the following sigmoidal function using PRISM 6 (GraphPad) 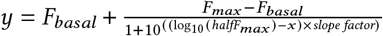 where *halfF*_*max*_ is the OD for which 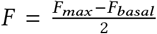. For cases with too weak fluorescence readings, the maximum fluorescence values *F*_*max*_ were extracted manually (typically the value read after 12 to 14 h of culture). Plotted averages and standard deviations were derived from at least two independent experiments, each one including several replicates per case (typically 3 clones). The data presented were normalized with respect to the value of the RMM pair, which served as positive control. Further experimental characterizations are detailed in Supplementary Methods.

### Design of nanobodies using de novo backbones

A structural model of the MR17 synthetic nanobody in interaction with the SARS-CoV-2 XBB.1.16 RBD was constructed by aligning the crystal structure of the MR17-RBD Wuhan complex (PDB 7C8W) [106] with the cryo-EM structure of the XBB.1.5 RBD-ACE2 complex (PDB 8VKP) [113] using PyMOL [127]. The K478 residue in the XBB.1.5 RBD was mutated to arginine using Custozyme+ to model the XBB.1.16 RBD. Missing residues in the MR17 structure and steric clashes at the MR17-RBD interface were resolved with MODELLER (v10.5) [128]. The resulting MR17-XBB.1.16 RBD complex was used as input to RFdiffusion [13] to generate multiple novel CDR backbones through its denoising neural network. Partial diffusion (github.com/RosettaCommons/RFdiffusion) was used in combination with the Radius Of Gyration (ROG) potential (*partial* _*T* ∈{1, 3, 5, 7, 9, 11, 13, 15}, *guide*_*scale* ∈ {0.25, 0.4, 0.55, 0.7, 0.85, 1.0, 1.15} with a cubic guide decay). The noising/denoising process was applied solely to the sybody structure and parts of its sequence were masked during the generation process (28 residues corresponding to the 3 CDRs of MR17). In this setting, an input structure is partially noised to serve as the initial input instead of starting with a completely random structure. Ten structures were generated for each combination of the given parameters by varying the random seed, resulting in a total of 560 structures.

ProteinMPNN was applied to these 560 backbones to design 10 sequences per structure, targeting a designable region of 36 residues, including the three CDRs and additional interface residues (Figure S26, Supporting Information). Sequence design used a sampling temperature of *T* = 0.1. Conformations of nanobody-RBD complexes were constructed using Custozyme+, relaxed using Rosetta FastRelax protocol [129] with the beta_nov16 score function [101]. Disulfide bonds between relevant cysteines were fixed (4 disulfide bonds on the RBD and 1 on the nanobody). Interactions of the top 100 complexes, ranked by Rosetta InterfaceAnalyzer ΔG [107], were further analyzed. From these, 14 nanobody designs showing the best Rosetta’s binding scores with the RBD, as well as having the best propensity to form hydrogen bonds and a high sequence diversity, were chosen for further characterization. 50 ns MD simulations were performed, using AMBER18 [130] with the all-atom ff14SB AMBER force field [131] under near-physiological conditions (*T* = 310 K, *P* = 1 bar, protonation states of titrable residues assigned at pH 6.5 using the pdb2pqr server [132], 150 mM NaCl concentration with the number of ions computed using the SPLIT method [133]). Details on MD simulations are provided in the Supporting Information. The 3 most stable complexes along MD simulations were then used as templates for EffieDes sequence design, using EffieNN (trained on multi-chain protein dataset with added std-dev 0.2 Å Gaussian noise on atomic coordinates) and the exact optimization algorithm of toulbar2. For each template, a total of 30 sequences were designed with, as above, a designable region containing 36 residues, and diversity constraints set to 2 amino acids between sequences. Six complexes were selected for MD simulations based on the same criteria as above: Rosetta binding scores with the RBD, propensity to form interfacial hydrogen bonds, and sequence diversity. From both ProteinMPNN and EffieDes pipelines, the 9 nanobodies (Table S13, Supporting Information) exhibiting the strongest interactions with RBD XBB.1.16, based on MM/GBSA and MM/PBSA calculations [108–112] over 50 ns MD trajectories, as well as showing the highest numbers of interfacial contacts and hydrogen bonds, were selected for experimental characterization. These nanobodies were named NbRM-P or NbRM-E (for **N**ano**b**odies generated with **R**Fdiffusion from the **M**R17 structure with a sequence predicted by **P**roteinMPNN or **E**ffieDes).

### Experimental characterization of nanobodies interaction with SARS-CoV-2 RBDs

RBDs of the SARS-CoV-2 spike protein (strains XBB.1.16 and Delta) were expressed in EXPI-293F cells (A14527) (Thermofisher), in fusion with a N-ter signal peptide and a C-ter His-tag, using the eukaryotic pYD11 expression plasmid under CMV promoter control.

Genes encoding nanobodies were cloned into pET24 expression vector, enabling expression of a C-terminal His-tagged protein. Plasmids were transformed into *E. coli* BL21(DE3) cells. For each nanobody, a 100 mL overnight preculture was prepared from a single colony in 2YT (or LB) medium supplemented with 50 µg mL^−1^kanamycin and incubated at 37 ^°^C with shaking at 200 rpm. Small-scale cultures of 12 mL LB medium were inoculated and incubated at 37 ^°^C. When the optical density at 600 nm reached 0.6, protein expression was induced with 1 mM IPTG. Cultures were incubated for an additional 5 h at 37 ^°^C under agitation before harvesting. Cells were then collected by centrifugation (5,000 × g, 10 min, 4 ^°^C) and pellets were resuspended in 2 mL of lysis buffer (50 mM Tris-HCl pH 8.0, 300 mM NaCl, 1 mg mL^−1^ lysozyme, 5 µg mL^−1^ DNase I, 5 mM MgCl_2_). Cell disruption was achieved by three freeze–thaw cycles (liquid nitrogen followed by 37 ^°^C water bath). Lysates were clarified by centrifugation (15,000 g, 15 min, 4 ^°^C), and the soluble fractions were collected for western blot and BLI analyses (see Supporting Information for more details and Figure S20-S21). Nanobodies and RBD proteins were both purified by affinity of their His-tag by IMAC on a nickel-charged resin, followed by size-exclusion chromatography using a Superdex 75 16/60 column (see Supporting Information and Figures S22-S23 for production and purification details).

Biolayer interferometry (BLI) was used to measure nanobodies interaction with the RBD. It is a label-free optical biosensing technique that measures biomolecular interactions in real time by monitoring wavelength shifts in the reflected light spectrum of a sensor surface during analyte binding. BLI experiments were performed on an Octet system (Octet RED96) (FortéBio, CA). A black bottom 96-well microplate (Greiner Bio-One # 655209) was filled with 200 µL of solution (nanobodies in PBS buffer) and agitated at 1,000 rpm, at 25 ^°^C. Tips were hydrated in PBS buffer for 20 min at room temperature prior to further experiments. Biotinylated SARS-CoV-2 RBDs (5 µg mL^−1^) were loaded on streptavidin SA (18-0009) biosensors (Pall ForteBio) for 1 min. After a baseline step in assay buffer (PBS [pH 7.4], 0.1% bovine serum albumin, 0.02% Tween 20) and a quenching step in 5 µg mL^−1^ biocytin, RBD-loaded sensors were dipped into known concentrations of NbRM-E1 nanobody (or total cell extract for interaction screening) for an association phase during 200-300 s. The sensors were then dipped in assay buffer for a dissociation step during 500 s. The association and dissociation curves were globally fitted to a 1:1 binding model. K_D_ was determined by fitting binding curves with the “association then dissociation” equation in the FortéBio Data analysis software version 7.1, resulting in K_D NbRM-E1_ = 64 nM.

## 5 Acknowledgments

M.D. and D.D. contributed equally to this work, and both have the right to place their name first. The authors thank Romain Gambardella for his suggestions on the neural architecture, and Justas Dauparas for sharing the multi-chain training dataset of ProteinMPNN. They are grateful to Shunsuke Tagami and Sota Yagi for valuable feedback on RNA polymerase. They also thank Isabelle Imbert and Juan Cortés for insightful discussions on nanobody design and characterization, as well as Jérémy Esque for discussions on BMC-H protein modeling. The authors acknowledge the PICT-ICEO platform for providing access to protein purification equipment. PICT-ICEO is a member of IBISBA-FR (https://doi.org/10.15454/08BX-VJ91), the French node of the European research infrastructure IBISBA (www.ibisba.eu). This work was supported by the French National Research Agency (ANR) under grant agreements ANR-19-CE09-0032-02, ANR-22-CE45-0025-01, ANR-19-PIA3-0004, the EUR BioEco (grant ANR-18-EURE-0021), and the ANITI AI cluster (ANR-23-IACL-0002). It benefited from high-performance computing (HPC) resources provided by CALMIP (allocations 2022-P21025, 2022-P22014, 2023, and 2024-P23015) and Jean-Zay GENCI-IDRIS (allocation 2022-AD011013779).

